# Mathematical Analysis of Robustness of Oscillations in Models of the Mammalian Circadian Clock

**DOI:** 10.1101/2020.09.15.297648

**Authors:** Benjamin Heidebrecht, Jing Chen, John J. Tyson

## Abstract

A wide variety of organisms possess endogenous circadian rhythms (~24 h period), which coordinate many physiological functions with the day-night cycle. These rhythms are mediated by a molecular mechanism based on transcription-translation feedback. A number of mathematical models have been developed to study features of the circadian clock in a variety of organisms. In this paper, we use bifurcation theory to explore properties of mathematical models based on Kim & Forger’s interpretation of the circadian clock in mammals. Their models are based on a simple negative feedback (SNF) loop between a regulatory protein (PER) and its transcriptional activator (BMAL). In their model, PER binds to BMAL to form a stoichiometric complex (PER:BMAL) that is inactive as a transcription factor. However, for oscillations to occur in the SNF model, the dissociation constant of the PER:BMAL complex, *K*_d_, must be smaller than 10^−3^ nM, orders of magnitude below the limit set by the biophysics of protein binding. We have relaxed this constraint by introducing two modifications to Kim & Forger’s SNF model: (1) replacing the first-order rate law for degradation of PER in the nucleus by a Michaelis-Menten rate law, and (2) introducing a multistep reaction chain for posttranslational modifications of PER. These modifications significantly increase the robustness of oscillations, and increase the maximum allowable *K*_d_ to more reasonable values, 1—100 nM. In a third modification, we considered alternative rate laws for gene transcription to resolve an unrealistically large rate of *PER* transcription at very low levels of BMAL transcription factor. Additionally, we studied Kim & Forger’s extensions of the SNF model to include a second negative feedback loop (involving REV-ERB) and a supplementary positive feedback loop (involving ROR). We found that the supplementary positive feedback loop—but not the supplementary negative feedback loop— provides additional robustness to the clock model.

**AUTHOR SUMMARY:** The circadian rhythm aligns bodily functions to the day/night cycle and is important for our health. The rhythm originates from an intracellular, molecular clock mechanism that mediates rhythmic gene expression. It is long understood that transcriptional negative feedback with sufficient time delay is key to generating circadian oscillations. However, some of the most widely cited mathematical models for the circadian clock suffer from problems of parameter “fragilities”. That is, sustained oscillations are possible only for physically unrealistic parameter values. A recent model by Kim and Forger nicely incorporates the inhibitory binding of PER, a key clock protein, to its transcription activator BMAL, but oscillations in their model require a binding affinity between PER and BMAL that is orders of magnitude lower than the physical limit of protein-protein binding. To rectify this problem, we make several physiologically credible modifications to the Kim-Forger model, which allow oscillations to occur with realistic binding affinity. The modified model is further extended to explore the potential roles of supplementary feedback loops in the mammalian clock mechanism. Ultimately, accurate models of the circadian clock will provide predictive tools for chronotherapy and chrono-pharmacology studies.

## INTRODUCTION

Most organisms experience perpetual day/night cycles and need to synchronize their physiological functions with this potent external driving rhythm of light and temperature (1). Endogenous circadian rhythms meet this demand. Across animals, plants, fungi and cyanobacteria, nearly all circadian rhythms share four major physiological characteristics (2). First, circadian rhythms are autonomous, meaning their timekeeping persists in the absence of external light/dark cues. Second, circadian rhythms are temperature-compensated, meaning that the autonomous cycle’s period is quite resistant to temperature variations over a physiological range (3–5). Third, circadian rhythms can be entrained to light/dark cycles considerably different from 24 hours (6); e.g., mammals can entrain to light/dark periods from 20.5 to 27.5 h (7). Fourth, the autonomous circadian rhythm can be phase-shifted by brief light exposures or temperature pulses. These perturbations cause a permanent shift in the phase of the oscillation without changing its period. This property underlies the familiar discomfort of ‘jet lag’ experienced by transcontinental travelers (8).

The circadian rhythms are generated by molecular clock mechanisms that generate oscillations of ~24h period through transcriptional-translational feedback (1, 9, 10). Although the genes and proteins constituting the circadian clocks in animals, plants and fungi are quite different, their essential interactions are remarkably similar. In all cases, the clock mechanism features a ‘core’ negative feedback loop: *A activates B activates C inhibits A*. In mammals, this loop consists of transcriptional regulation involving six genes: *PER1/2*, *CRY1/2*, *BMAL1*, and *CLOCK* (1, 9–11). For convenience, in this work we drop the distinction between the homologous pairs of proteins PER1/2 and CRY1/2. In this mechanism (Figure 1), the heterodimeric transcription factor BMAL:CLOCK activates *PER* transcription. *PER* mRNA is then translated in the cytoplasm, where PER protein binds with CRY and enters the nucleus. PER:CRY then binds with BMAL:CLOCK to block its activation of *PER* transcription. PER:CRY’s cycle of production, nuclear entry, auto-inhibition, and subsequent degradation is widely acknowledged to be the source of circadian rhythmicity (12). BMAL:CLOCK contributes to genome-wide rhythmic transcription; for instance, in mouse liver, BMAL:CLOCK targets the enhancers of ~1,300 genes, out of which ~26% are rhythmically transcribed (13). Additional transcriptional regulation comes from REV-ERB and ROR proteins. BMAL:CLOCK activates the expression of their genes, and subsequently, cytoplasmic ROR and REV-ERB proteins are translocated into the nucleus where they compete for repression (by REV-ERB) or activation (by ROR) of *BMAL1* transcription (Figure 1) (1, 9–11). Ultimately, 3―15% of genes are rhythmically expressed in any tissue at any time (14), and the clock transcription factors play important roles in this program (13, 15).

**Figure 1.**
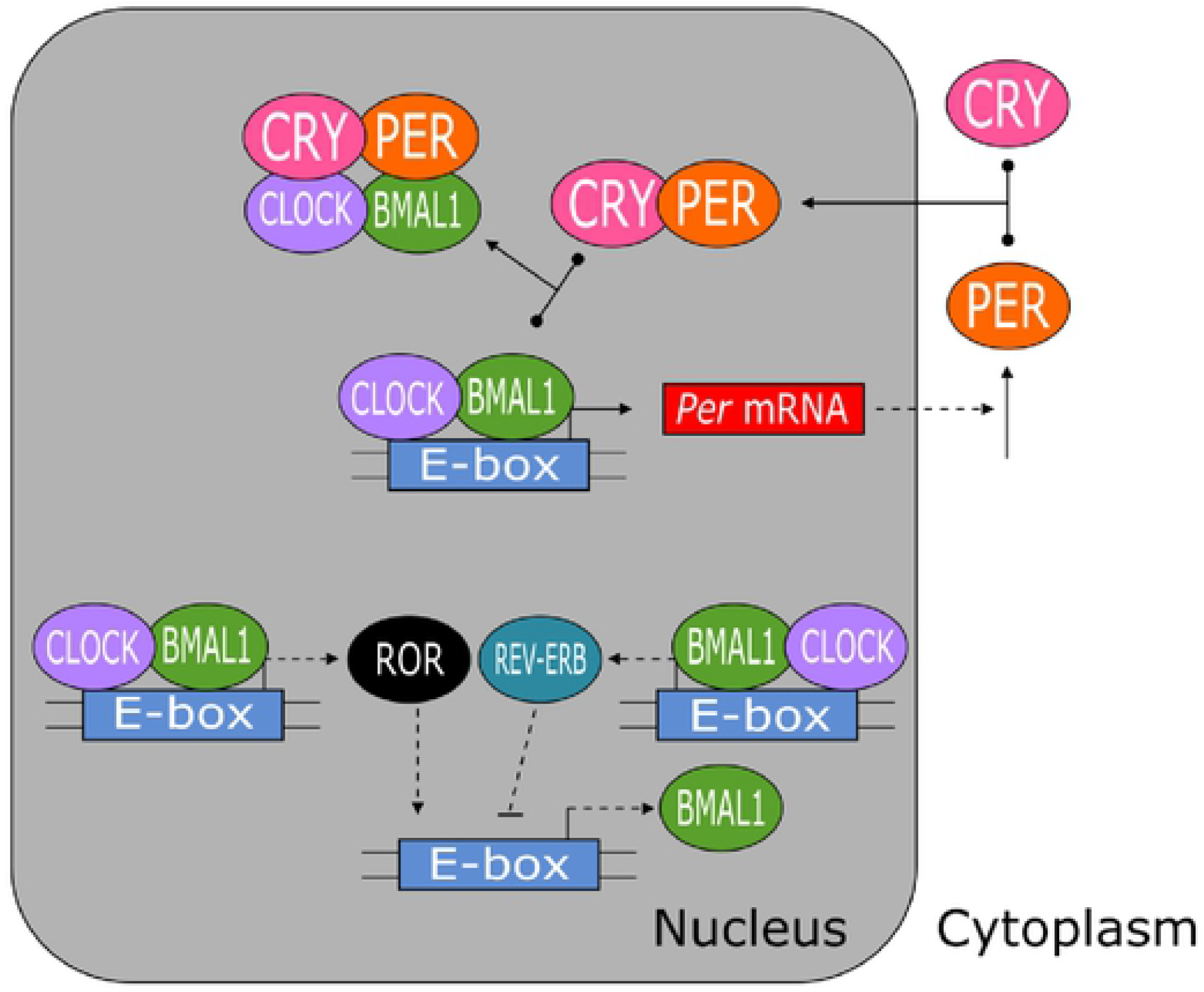
Three major feedbackloops regulate the mammalian circadian clock. The core negative feedback loop is between PER:CRY and BMALl:CLOCK. The two sources of additional feedback are negative feedback between REV ERB and BMALl and positive feedback between ROR and BMALl. PERl/2, CRYl/2, and RORα/β are simpl fied to PER, CRY, and ROR, respect vely. Solid lines indicate chemical reactions; the T shaped reactions indicate reversible binding of proteins to form multicomponent complexes. Dashed lines indicate regulatory s gnals (positive regulation = barbed arrow, and negative regulation = blunt arrow).

Over the past 50 years, many people have proposed mathematical models of circadian rhythms (3, 12, 16–20). In 1965, Brian Goodwin proposed a model of periodic enzyme synthesis based on negative feedback on gene expression (21, 22). Goodwin was not attending to circadian rhythms, at the time, because nothing was known then about the negative feedback of PER on its own synthesis. But his model was picked up later by Peter Ruoff (5, 23–25) to explain the characteristic features of circadian rhythms. Recently, the core negative feedback loop of Goodwin’s model was extended with other feedback loops (as in Figure 1) to create more comprehensive and realistic models of circadian rhythms (26–28). One particularly interesting modification to Goodwin’s model was made by Jae Kyoung Kim and Daniel Forger (27), who replaced Goodwin’s view of negative feedback by cooperative multimeric binding of a generic ‘repressor’ to a gene promoter with their own model of stoichiometric binding of PER:CRY (a repressor) to BMAL:CLOCK (an activator) of gene expression. Some characteristic features of the two models were compared in (29). While all of these models have much to commend, they suffer from some technical problems (parameter ‘fragilities’) that limit their appeal. In the following section, we outline these difficulties, in order to frame our proposals for more robust and realistic mathematical models of circadian clocks. Then, in the ‘Results and Discussion’ section, we present our results in detail.

## PRELIMINARY MODELING ISSUES

### Goodwin’s Model

To account for observations of periodic enzyme synthesis in bacteria (30), Brian Goodwin (21, 22) presented the following model for the periodic synthesis of a protein Y from its mRNA X, where mRNA synthesis is inhibited by a repressor Z:

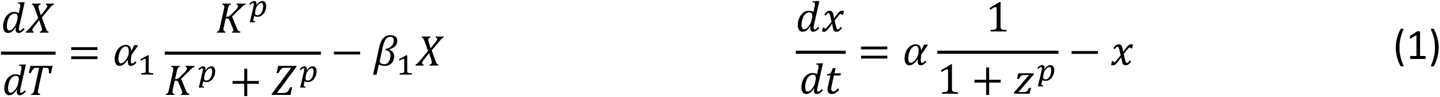

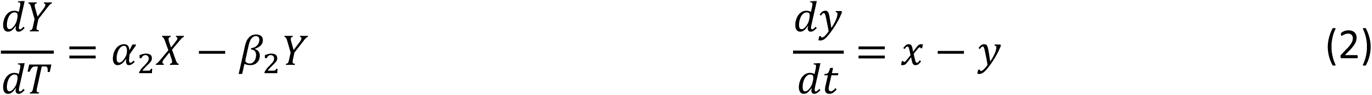

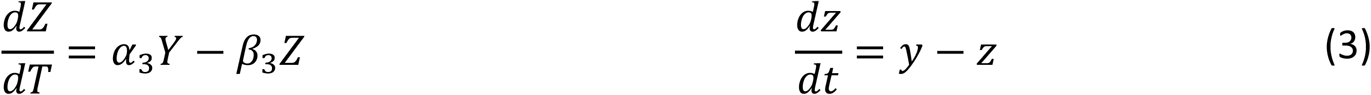

In Eq. (1), the factor 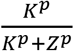 is the probability that the promoter region of the gene encoding X is <u>not</u> bound to Z, its inhibitor, and *α*1 is the maximum rate of synthesis of X by the gene. The other terms in these equations correspond to first-order rate laws for production and removal of X, Y and Z. In this tableau, Goodwin’s equations are written in two equivalent forms. On the left, the equations are written in terms of the original dimensional variables: concentrations *X*, *Y* and *Z* (nM) and time *T* (h); on the right, in terms of the ‘dimensionless’ variables 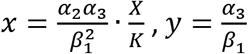. 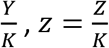, dimensionless time *t* = *β*_1_ *T*, and a dimensionless parameter 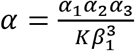. In deriving these dimensionless equations, we have assumed (as have all authors in the past) that *β*_1_ = *β*_2_ = *β*_3_, which serves to maximize the oscillatory potential of the model (31, 32). In Goodwin’s version of a three-component negative-feedback loop, the repression of gene transcription by Z is modeled by a Hill function with exponent *p*. Underlying this function is the supposition that the gene encoding X is turned off when *p* molecules of Z bind cooperatively to its promoter region (or, equivalently, when *p* molecules of Z bind cooperatively to an activator of gene transcription and shut it off).

#### A problem with Goodwin’s model

J.S. Griffith (33) was first to point out that Goodwin’s equations (1)-(3) admit oscillatory solutions only if *p* > sec^3^(*π*/3) = 8, a very restrictive condition, because in experimental studies it is rare that more than 3 or 4 proteins bind cooperatively to DNA regulatory sequences (34). This condition becomes even more restrictive if *β*_1_ ≠ *β*_2_ ≠ *β*_3_ (31).

#### One solution: a longer feedback loop

The restriction *p* > 8 can be ameliorated by lengthening the feedback loop: if *n* = number of variables in the feedback loop, then the condition becomes *p* > sec^*n*^(*π/n*). For example, for *n* = 8, the condition is *p* > 1.88. Longer loops (*n* > 3) correspond to inserting more than one intermediate (say, Y_0_, Y_1_, …, Y_*n*−3_) between X (mRNA) and Z (feedback component). This is quite reasonable, considering that PER protein has multiple phosphorylation sites (35). Each intermediate, Y_*j*_, then denotes cytoplasmic PER phosphorylated on *j* sites, *j* = 0, 1, …, *n*−3. Eventually, the fully phosphorylated form, Y_*n*−3_, is transported into the nucleus and becomes Z. In this case, Goodwin’s dimensionless differential equations become Eqs. (4)-(7).

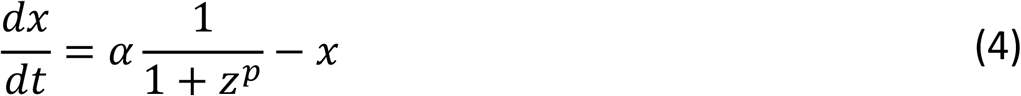

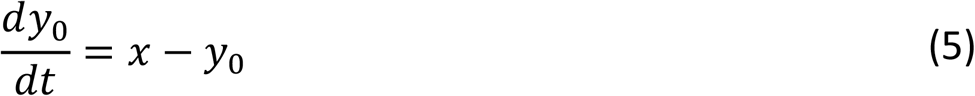

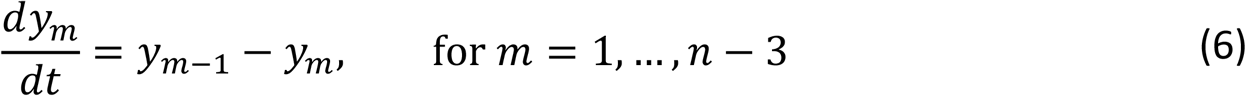

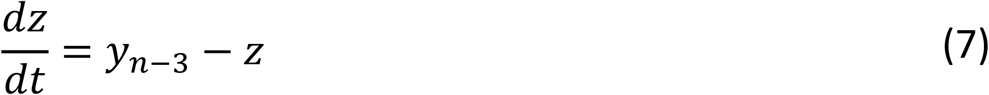

Exactly the same equations can be derived by assuming a distributed time lag between X and Z (36) and introducing the new variables

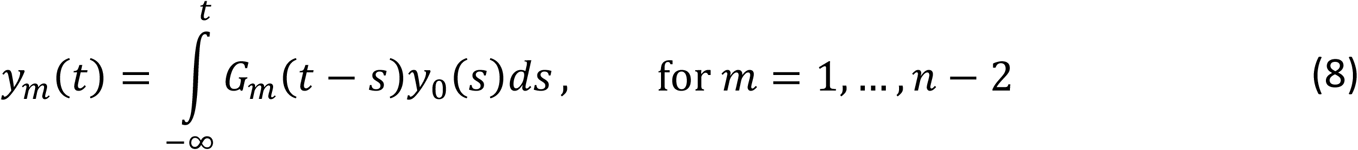

where *y*_0_(*t*) is the dimensionless form of *Y*_0_(*T*), and *G*_*m*_(*u*) = *u*^*m*−1^*e*^−*u*^/(*m* − 1)! for integer values of *m*. From this definition we can show that

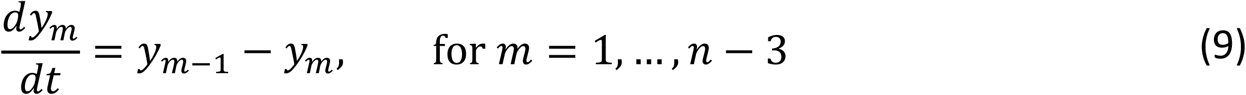

which is exactly Eq. (6) above.

#### A second solution: Michaelis-Menten degradation of Z

In 1982 Bliss, Painter and Marr (37) proposed to replace the first-order degradation of Z, −*β*_3_*Z*, by a Michaelis-Menten rate law, −*β*_3_*Z*/(*K*_m_ + *Z*), where *K*_m_ is the ‘Michaelis’ constant of the enzyme-catalyzed reaction and *β*_3_ is the ‘*V*_max_’ of the reaction. With this change, the Goodwin model can exhibit limit cycle oscillations even for *p* = 1 (37). The substitution of Michaelis-Menten rate laws for the first-order kinetic terms in Eqs. (1)-(3) has been exploited by many authors (17, 38, 39) to increase the robustness of their models of circadian rhythms.

### Kim & Forger’s Model

In 2012, Kim and Forger (27) presented a slightly different model of the negative feedback loop controlling mammalian circadian rhythms (Figure 2a). In dimensionless form, the Kim-Forger (KF) ODEs are:

**Figure 2.**
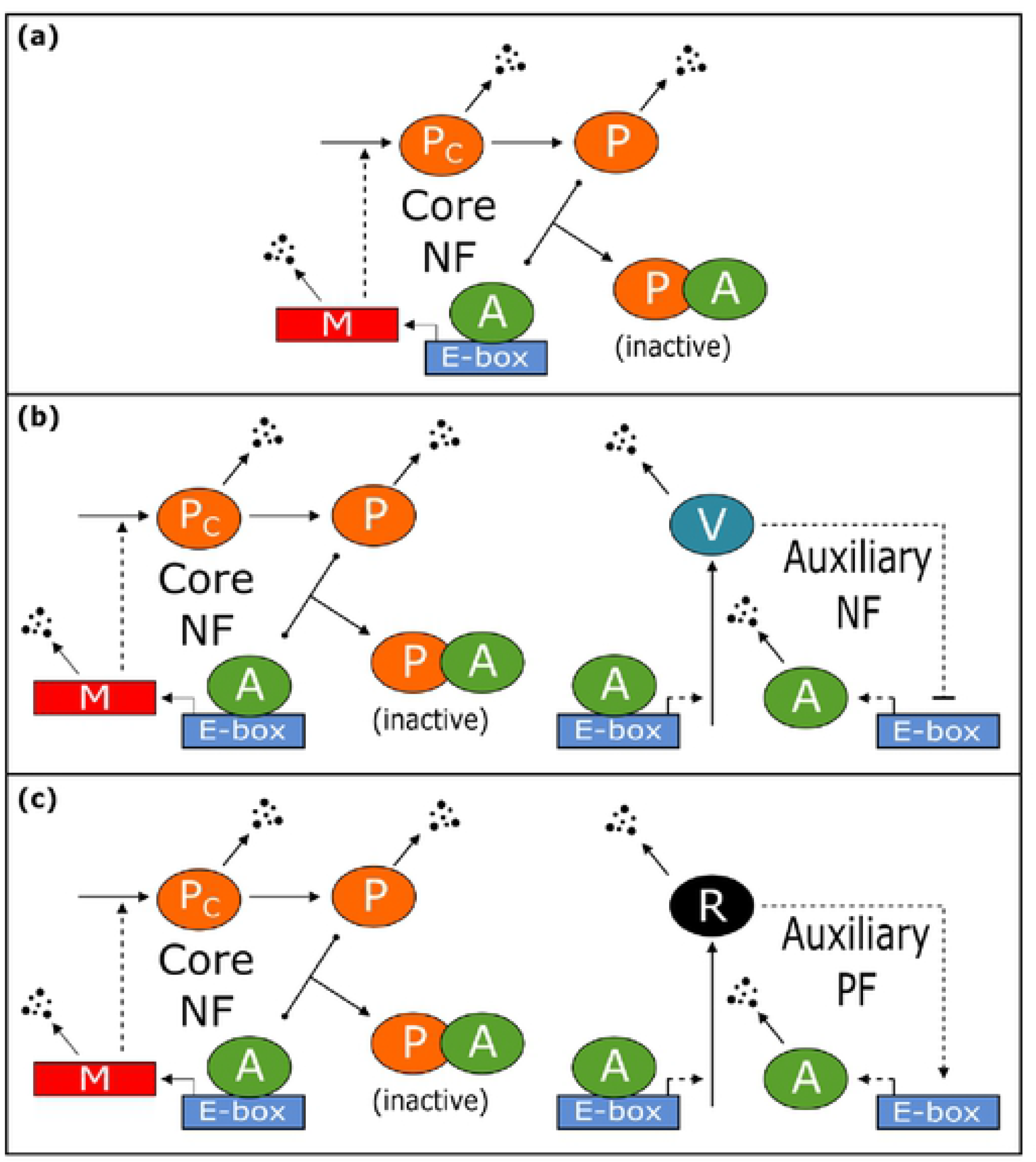
Wiring diagrams for the three Kim-Forger models: SNF **(a)**, NNF **(b)**, and PNF **(c)**. To simplify the models, several molecu lar species that do not contribute significantly to the feedback loops are not explicitly represented. For example, in the SNF loop, CLOCK and CRYare not shown. In the NNF and PNF loops, mRNAs encoding REV-ERB, ROR and BMAL are not shown, nor are the cytoplasmic forms of these proteins. Solidand dashed lines indicate reactions and regulations, as in Figure 1.

#### Kim-Forger SNF Model

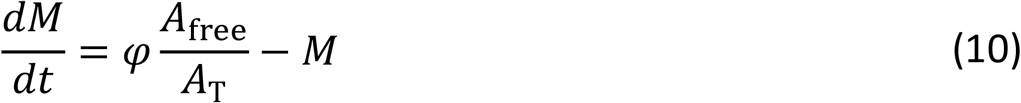

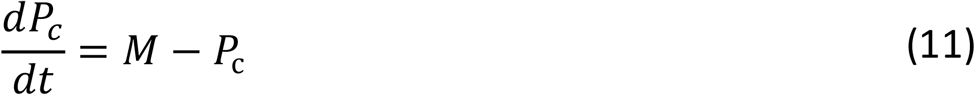

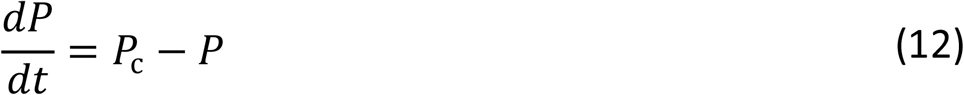

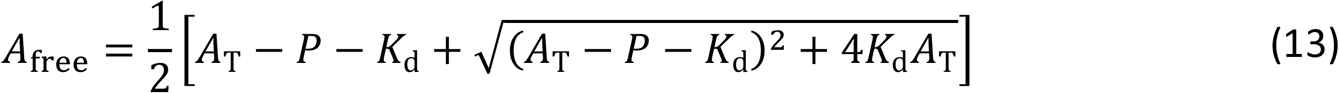

SNF stands for ‘simple negative feedback loop’ (i.e., the negative feedback loop involving PER:CRY inhibition of BMAL:CLOCK). As originally written, the KF model has three dynamical variables: *M* = [*PER* mRNA], *P*_c_ = [PER protein in the cytoplasm], *P* = [PER protein in the nucleus] (i.e., PER:CRY in the nucleus). The BMAL:CLOCK transcription factor is denoted by A; *A*_T_ is the total concentration of BMAL:CLOCK in the nucleus, and *A*_free_ is the concentration of ‘free’ BMAL:CLOCK in the nucleus. The factor *A*_free_/*A*_T_ is the probability that BMAL:CLOCK is <u>not</u> bound to its repressor, PER:CRY. By expressing the rate of transcription of *PER* mRNA to be proportional to *A*_free_/*A*_T_, Kim & Forger are implicitly assuming that the total number of BMAL:CLOCK dimers is large enough to saturate the E-boxes on the *PER* genes, and that PER:CRY binds equally well to BMAL:CLOCK dimers that are either bound or not bound to an E-box (Supplementary Materials, ‘Deriving the rate laws for *PER* transcription’). The parameter *φ* represents the gene dosage of *PER*, relative to homozygous diploid, *φ* = 1. For cells over- (or under-) expressing the *PER* gene, *φ* > 1 (or *φ* < 1). The expression for *A*_free_ in Eq. (13) is derived by solving the condition for equilibrium binding of BMAL:CLOCK (A) and PER:CRY (P) to form an inactive complex (C); namely, *K*_d_*C* = *A*_free_*P*_free_, where *A*_free_ = *A*_T_ − *C* and *P*_free_ = *P* − *C*.

The KF equations (10)-(13) are written in dimensionless form; in particular, *t* = *β* × time, where *β* is the first-order rate constant (h^−1^) for removal of all three species in Eqs. (10)-(12). Furthermore, *P*, *K*_d_, *A*_T_ and *A*_free_ are all scaled by the same factor, *P*\* ≈ 25 nM (Supplementary Materials, ‘Estimation of the scaling factor *P*\*’).

Notice that, in the KF model, nonlinearity in the transcription term is due to tight stoichiometric binding between PER:CRY and BMAL:CLOCK, not (as in Goodwin’s equations) to cooperative participation of nuclear PER in the regulation of *PER* gene expression. Consequently, the KF model circumvents the unreasonable cooperativity constraint (*p* > 8) of Goodwin’s model. (Don’t confuse the Hill exponent, *p*, in Goodwin’s model with the concentration of nuclear PER, *P*, in the KF model.)

In addition to the SNF model, Kim & Forger proposed two extended models, in which the core negative feedback loop involving PER and BMAL is supplemented with (either) an additional <u>negative</u> feedback from REV-ERB on transcription of the *BMAL1* gene (called the NNF model, Figure 2b) (or) an additional <u>positive</u> feedback from ROR on transcription of the *BMAL1* gene (called the PNF model, Figure 2c). Both the NNF and PNF models include the ODEs of the core SNF model.

#### Kim-Forger NNF Model

Equations (10)-(13), plus

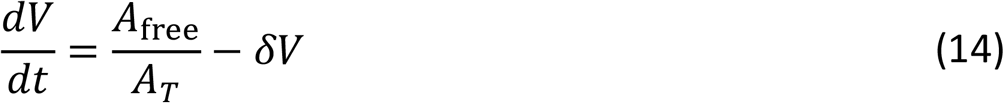

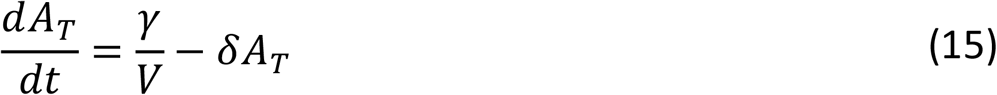

where *V* is the (scaled) concentration of REV-ERB, *γ* is a rate constant for *BMAL1* transcription, and *δ* is a rate constant that sets the time scale for the feedback loop.

#### Kim-Forger PNF Model

Equations (10)-(13), plus

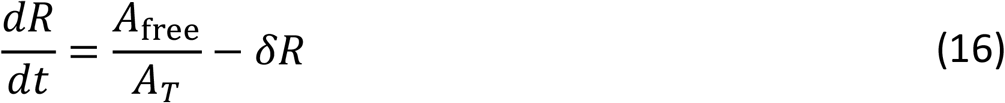

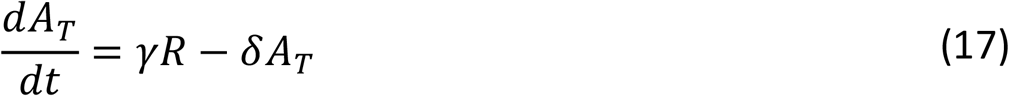

where *R* is the (scaled) concentration of ROR, and *γ* and *δ* are defined similarly as in the NNF equations.

In Tables S1 and S2 we provide definitions of all the variables and kinetic constants in these models.

While the KF models appear to oscillate robustly and avoid Goodwin’s unrealistic constraint (*p* > 8), the core SNF model seems to have an unreasonable constraint of its own; namely, the maximum permissible value of the dissociation constant of the nuclear PER:BMAL complex is *K*_d_ = 10^−4^, which is orders of magnitude too small for protein-protein binding. To see why, we convert KF’s dimensionless *K*_d_ into its dimensional form, *K*_d_ < 10^−3^ nM, by multiplying by *P*\* ≈ 10 nM (this is an order-of-magnitude estimate). Because the time constant for dissociation of the PER:BMAL complex is expected to be on the order of minutes (i.e., *k*_unbind_ ≈ 0.01 s^−1^), the binding constant for the complex would have to be 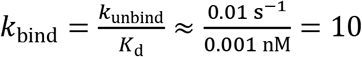 nM^−1^s^−1^ = 10^10^ M^−1^ s^−1^. However, protein-protein binding rate constants are typically on the order of 10^6^ M^−1^s^−1^ (40). So, a physically realistic value for the dissociation constant of the PER:BMAL complex is *K*_d_ ≈ 10 nM, or in dimensionless terms, *K*_d_ ≈ 1. Even if the binding is especially tight, *K*_d_ should be at least on the order of 0.1. In fact, Fribourgh et al. (41), who recently studied the docking of PER2:CRY1/2 to the core PAS domain of BMAL:CLOCK, measured *K*_d_ ≈ 400 nM, or in dimensionless terms, *K*_d_ ≈ 40. This estimate may easily be off by a factor of 10, because the measurements were made *in vitro* with purified protein domains (not with full length proteins). Nonetheless, comparing our theoretical estimate and this experimental result, we might conclude that *K*_d_ ≈ 0.1—10, in dimensionless terms.

Hence, KF’s dissociation constant is unrealistically small by several orders of magnitude. In this work we consider some realistic changes to the SNF model that will increase the robustness of its oscillations. In the process, we come up with some other surprising reassessments of the KF model.

## RESULTS AND DISCUSSION

### Longer Feedback Loop and Saturating PER Degradation Increase the Oscillatory Robustness of the Kim-Forger SNF Model

Our primary goal in modifying KF models is to alleviate the unreasonable constraint on *K*_d_, the dissociation constant of the PER:BMAL complex. To this end, we considered the same two changes that were made to Goodwin’s model to increase its robustness: first, increasing the number of dynamical species in the PER-BMAL negative feedback loop, and second, introducing a Michaelis-Menten rate law for the degradation of nuclear PER.

#### Longer feedback loop

Given that PER has 10-20 phosphorylation sites (35), it is reasonable to assume that cytoplasmic PER is phosphorylated sequentially on 1, …, *n*-3 sites before it is transported into the nucleus. Hence, we rewrite P_C_ as P_0_ and insert between Eq. (11) and Eq. (12) a sequence of linear ODEs for the intermediate (phosphorylated) states, P_1_, …, P_*n*-3_. This change lengthens the time between *PER* mRNA transcription and the negative feedback signal generated by nuclear PER and consequently increases the oscillatory potential of the negative feedback loop (42).

#### Saturating degradation of nuclear PER

PER is degraded by proteasomes after it is poly-ubiquitinated by the E3 ligase β-Trcp (43). Because the rate of this enzyme-mediated reaction likely saturates at large substrate concentration, it is reasonable to replace the linear kinetics for nuclear PER degradation in Eq. (12) by a Michaelis-Menten rate law (43),

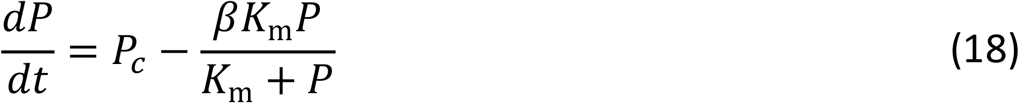

In Eq. (18), *K*_m_ is the Michaelis constant of the rate law and *βK*_m_ is the ‘*V*_max_’. This change also has the potential to increase the oscillatory robustness of the model. Intuitively, the upper limit to the rate of PER degradation introduced by the Michaelis-Menten rate law causes nuclear PER concentration to react sluggishly to changes in the rate of *PER* mRNA production, which is another sort of ‘lag’ in the negative feedback loop.

Both modifications indeed improve the robustness of oscillations in the SNF model. ‘Longer feedback loops’ (with no other modifications of the SNF model) can increase by a hundred fold the maximum *K*_d_ for oscillations, as shown in Figure 3a. This upper bound is still below the range of physical acceptability. If we introduce both ‘saturating degradation’ and an ‘*n*-component feedback loop’, then the maximum *K*_d_ for oscillations increases to a value of 1 (see Figure S1), but this happy result is suspect because the doubly-modified SNF model exhibits oscillations even as *A_T_* → 0 when *n* ≥ 4. This result is impossible because there can be no expression of the *PER* gene when BMAL concentration is zero. The problem, of course, is that the rate law for *PER* transcription (rate ∝ *A*_free_/*A*_T_) is valid only if BMAL saturates the *PER* E-boxes. To get around this restriction, we consider some other potential rate laws for *PER* gene transcription (see Supplementary Materials, ‘Deriving the rate laws for *PER* transcription’).

**Figure 3.**
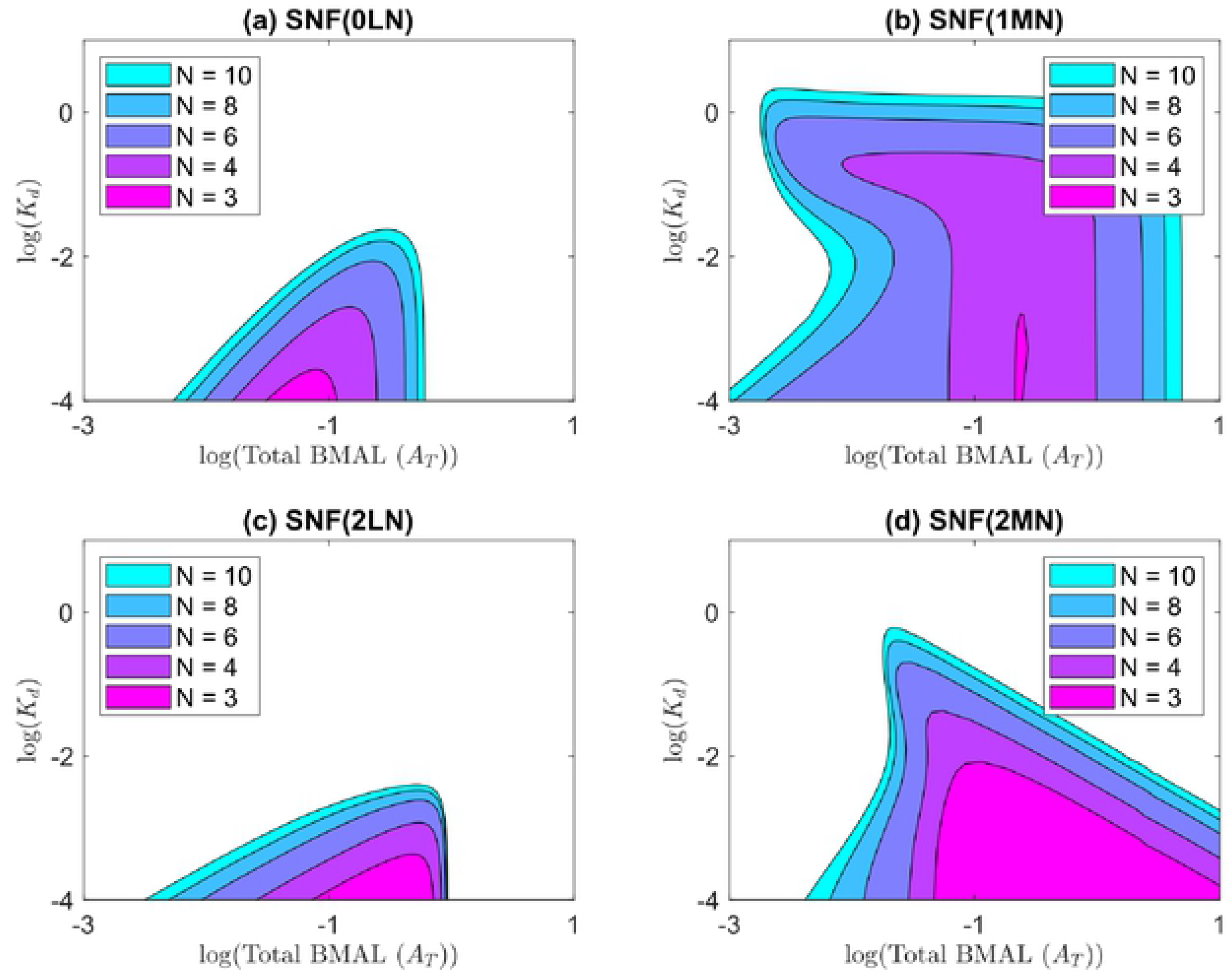
Two-parameter bifurcation diagrams, with respect to *K*_d_ and *A*_T_. **(a)** The original KF SNF(OLN) mode, with linear rate of degradation of nuclear PER. **(b)** SNF(lMN) model, with Michaelis-Menten degradation of nuclear PER. **(c)** SNF(2LN) model, with rate law 2 for *PER* mRNA transcription and linear degradation of nuclear PER. **(d)** SNF(2MN) model, with rate law 2 for *PER* mRNA transcription and Michaelis-Menten degradation of nuclear PER. Only model SNF(lMN) admits oscillations for reasonable values of *K*_d_ ≈ 1. The parameter values used in these calculations are provided in Table S3; in particular, *φ* = 1.

### Modifying the PER Transcription Rate Law Increases the Robustness of the SNF Model Still Further

We consider two slightly different rate laws to replace the KF expression for the rate of *PER* gene transcription:

<u>Rate Law 0</u>:

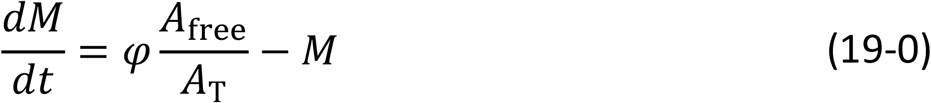

<u>Rate Law 1</u>:

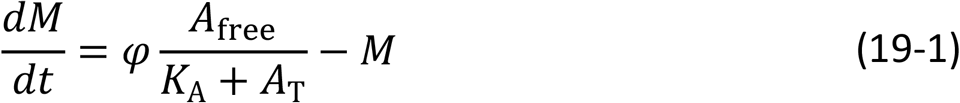

<u>Rate Law 2</u>:

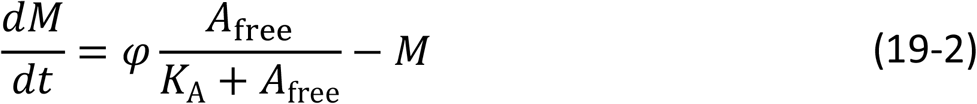

Rate law 0 is, of course, the original KF expression. When *A*_free_ = *A*_T_, the maximum rate of transcription is *φ*. (In our scaling of the differential equations, the maximum rate of transcription of *PER* mRNA = *φ* = 1 for WT homozygous diploid cells, and the rate increases or decreases with *φ* for cells over- or under-expressing *PER* mRNA, either by manipulating *PER* gene dosage or the strength of the *PER* gene promoter.) For rate laws 1 and 2, the maximum rate of transcription is 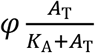, i.e., the maximum rate depends on how strongly BMAL:CLOCK binds to the E-box, characterized by the dissociation constant *K*_A_. Rate laws 1 and 2 also imply that the transcription rate is proportional to *A*_free_ (not *A*_free_ /*A*_T_) when *A*_T_ becomes small. These three rate laws represent different limiting cases of BMAL:CLOCK binding to E-boxes, as explained in detail in the Supplementary Materials, ‘Deriving the rate laws for *PER* transcription’. Recall that the KF rate law 0 applies to the case in which binding between PER:CRY and BMAL:CLOCK is independent of the binding between BMAL:CLOCK and E-box, and BMAL:CLOCK complexes saturate *PER* E-boxes. Rate law 1 is a slight modification of the KF rate law 0, relaxing the saturation of *PER* E-boxes by BMAL:CLOCK. Finally, rate law 2 is based on a different model of the regulation of *PER* transcription, where PER:CRY binds tightly to BMAL:CLOCK, and only the free form of BMAL:CLOCK binds to the E-box.

#### Modified Kim-Forger SNF equations

Taking all of the aforementioned changes into account, we have:

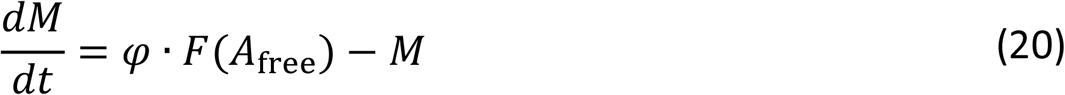

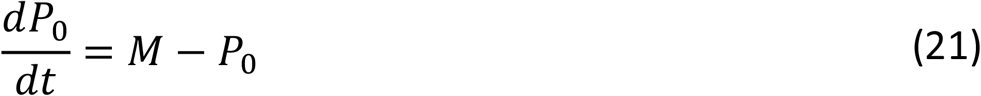

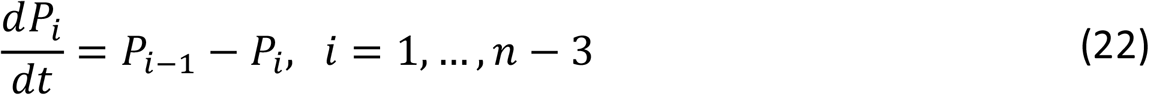

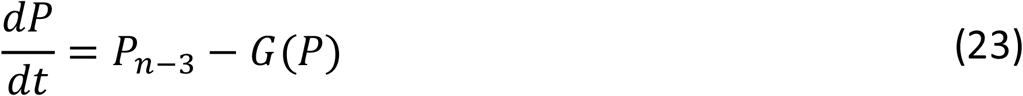

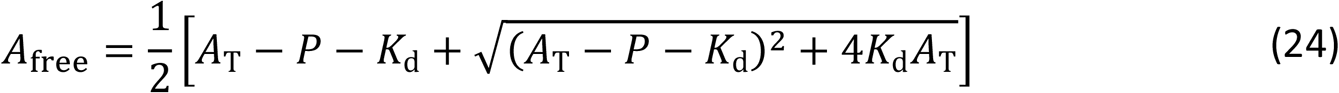

where 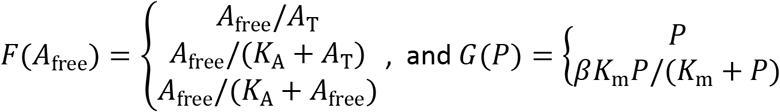.

To keep track of all these different versions, we introduce the notation SNF(TDN), where T denotes the *PER* transcription rate law (0, 1 or 2), D denotes the PER degradation rate law (L for ‘linear’ or M for Michaelian), and N denotes the number of species in the negative feedback loop (Supplementary Materials, ‘Model naming convention’). For example, the original KF SNF model is denoted SNF(0L3).

We have already shown the two-parameter bifurcation diagram (log *K*_d_ versus log *A*_T_) for SNF(0LN) in Figure 3a. The corresponding bifurcation diagram for SNF(1MN), Figure 3b, shows that our extension of the SNF model (with modified transcription rate and saturating degradation of nuclear PER) is more robust than the original KF model in that oscillations are now possible for *K*_d_ as large as 1. In Figures 3c and d, we display the matching bifurcation diagrams for SNF(2LN) and SNF(2MN), respectively. Again, we see that saturating degradation of nuclear PER is crucial for permitting oscillations for *K*_d_ > 0.01. Comparing Figure 3b and 3d, we see that transcription rate law 1 generates more robust oscillations than rate law 2 for *K*_d_ values larger than 0.1, but rate law 2 is more robust for N = 3.

### An Additional Positive Feedback Loop Involving ROR Increases the Domain of Oscillations

Next, we explore Kim & Forger’s NNF and PNF models (above), with modified rate laws for gene transcription. For the rates of transcription of *PER*, *REV-ERB* and *ROR* genes, we use Rate Law 2, Eq. (19–0), and for the transcriptional activation and repression of the *BMAL* gene by ROR (variable *R*) and REV-ERB (variable *V*), we replace the functions *γR* and *γ/V* (as originally proposed by Kim & Forger) by *A*_MAX_∙*R*/(*R*+1) and *A*_MAX_/(*V*+1), respectively. The new rate laws assume that *R* and *V* are scaled by the dissociation constants of these proteins with the promoter of the *BMAL* gene. These new rate laws remedy an issue in KF’s original PNF and NNF models, for which the rate of BMAL synthesis does not saturate as *R* →∞ or *V* → 0.

#### Modified Kim-Forger NNF model

Equations (20)-(24) with *F*(*A*_free_) = *A*_free_/(*K*_A_ + *A*_free_) plus

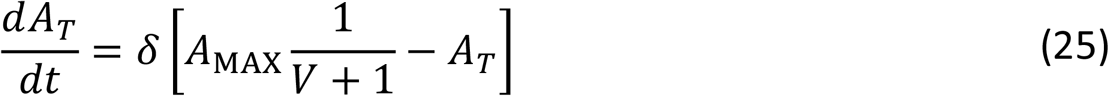

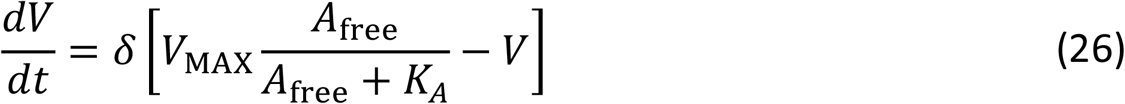

#### Modified Kim-Forger PNF model

Equations (20)-(24) with *F*(*A*_free_) = *A*_free_/(*K*_A_ + *A*_free_) plus

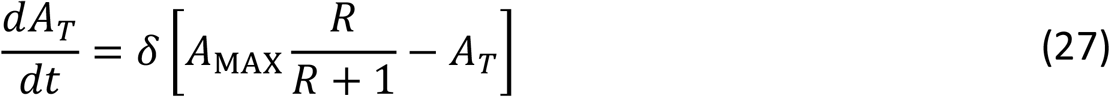

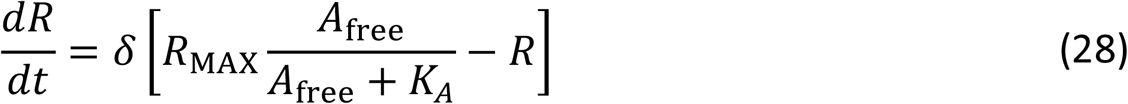

Finally, in wild-type SCN cells, nuclear REV-ERB and ROR compete for binding to the *BMAL* promoter. So, it is reasonable to consider a model that accounts for the effects of REV-ERB and ROR together. (This case was not considered by Kim and Forger.) In the new model (PNNF) the BMAL synthesis term accounts for competitive regulation by both REV-ERB and ROR, and also features a basal synthesis rate (*ε*) for the case when *R* and *V* are both zero. The PNNF model assumes that REV-ERB and ROR bind to the *BMAL* promoter with the same affinity.

#### PNNF equations

Equations (20)-(24) with *F*(*A*_free_) = *A*_free_/(*K*_A_ + *A*_free_) plus

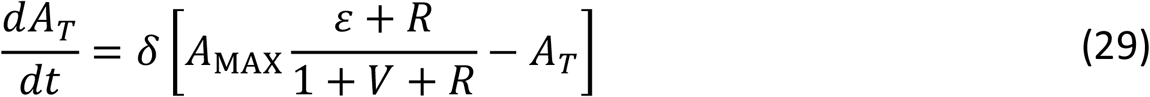

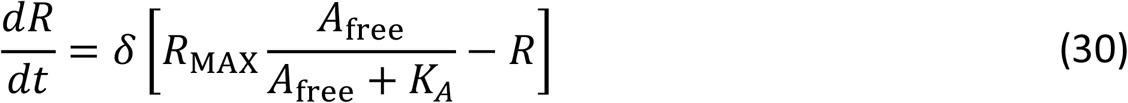

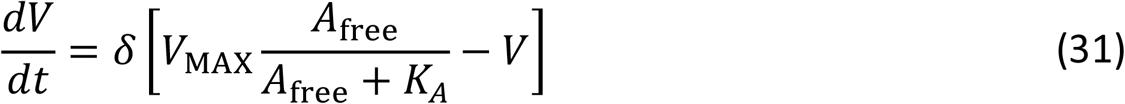

The original KF models (SNF, NNF, and PNF) were compared to each other by plotting the regions where each model oscillates with respect to fold changes in *BMAL* and *PER* transcription rates, Figure 4a, which is identical to Figure 5a in the Kim-Forger paper (27). The major conclusion from Kim & Forger’s calculations was that the additional negative feedback loop increases oscillatory robustness of the model while the additional positive feedback loop decreases oscillatory robustness.

**Figure 4.**
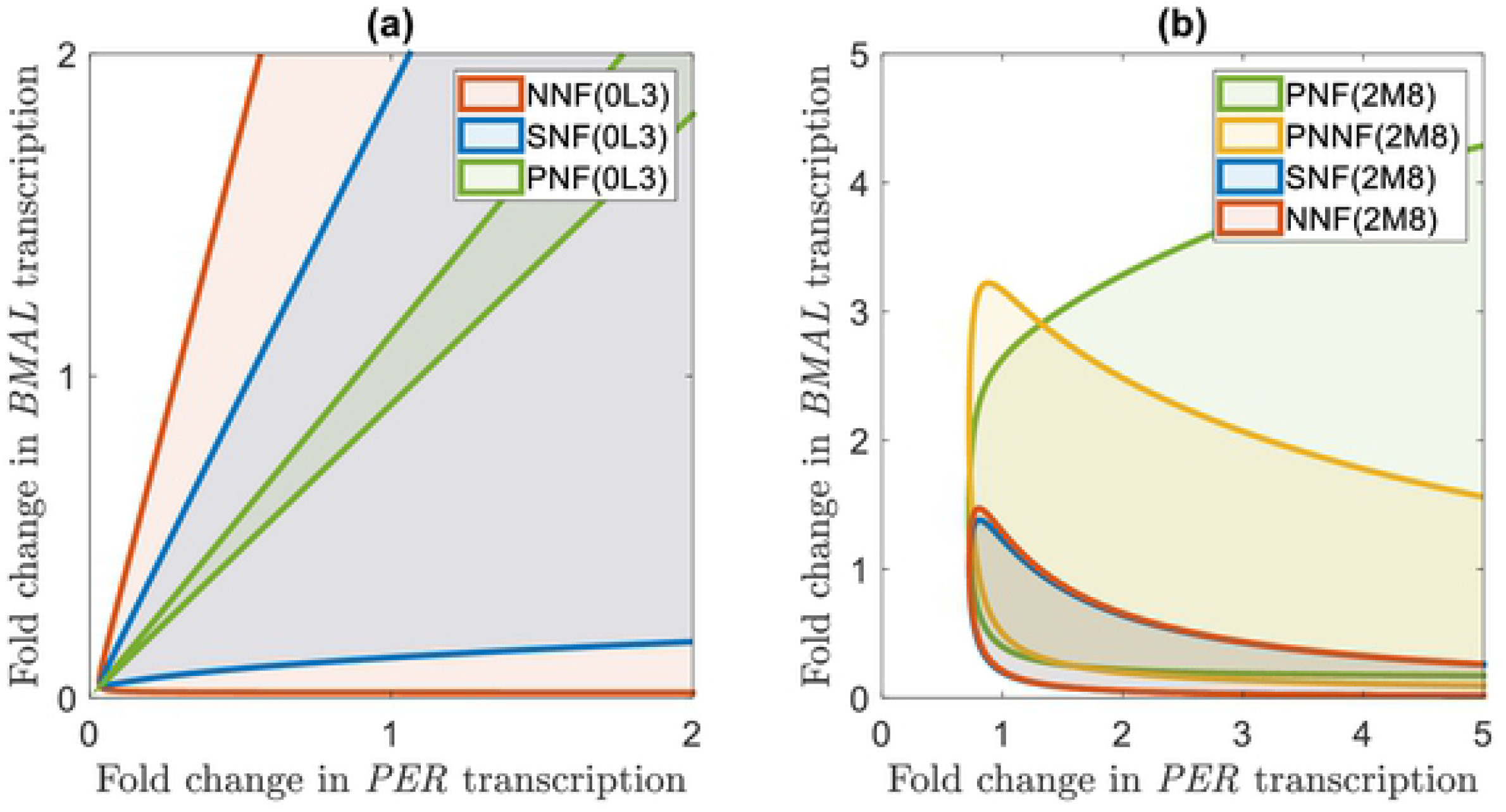
Two-parameter bifurcation curves with respect to fold changes in *BMAL* and *PER* transcription rates. **(a)** Comparison of oscillatory robustness for the unmodified KF models: SNF(OL3), NNF(OL3) and PNF(OL3). The *BMAL* transcription rate is varied with parameter *A_T_* (for SNF) or *γ* (for NNF and PNF). *K*_d_ = 10^−5^ in panel a (to allow oscillations in the unmodified KF models). **(b)** Comparison of oscillatory robustness for the modified KF models: SNF(2M8), NNF(2M8), PNF(2M8), PNNF(2M8). The *BMAL* transcription rate is varied with parameter *AT* (for SNF) or *A*_MAX_ (for NNF, PNF and PNNF). *K*_d_ = 10^−1^ in panel b. The *PER* transcription rate is varied with *φ* for all models. The other parameter values used in these calculations are provided in Table S4.

**Figure 5.**
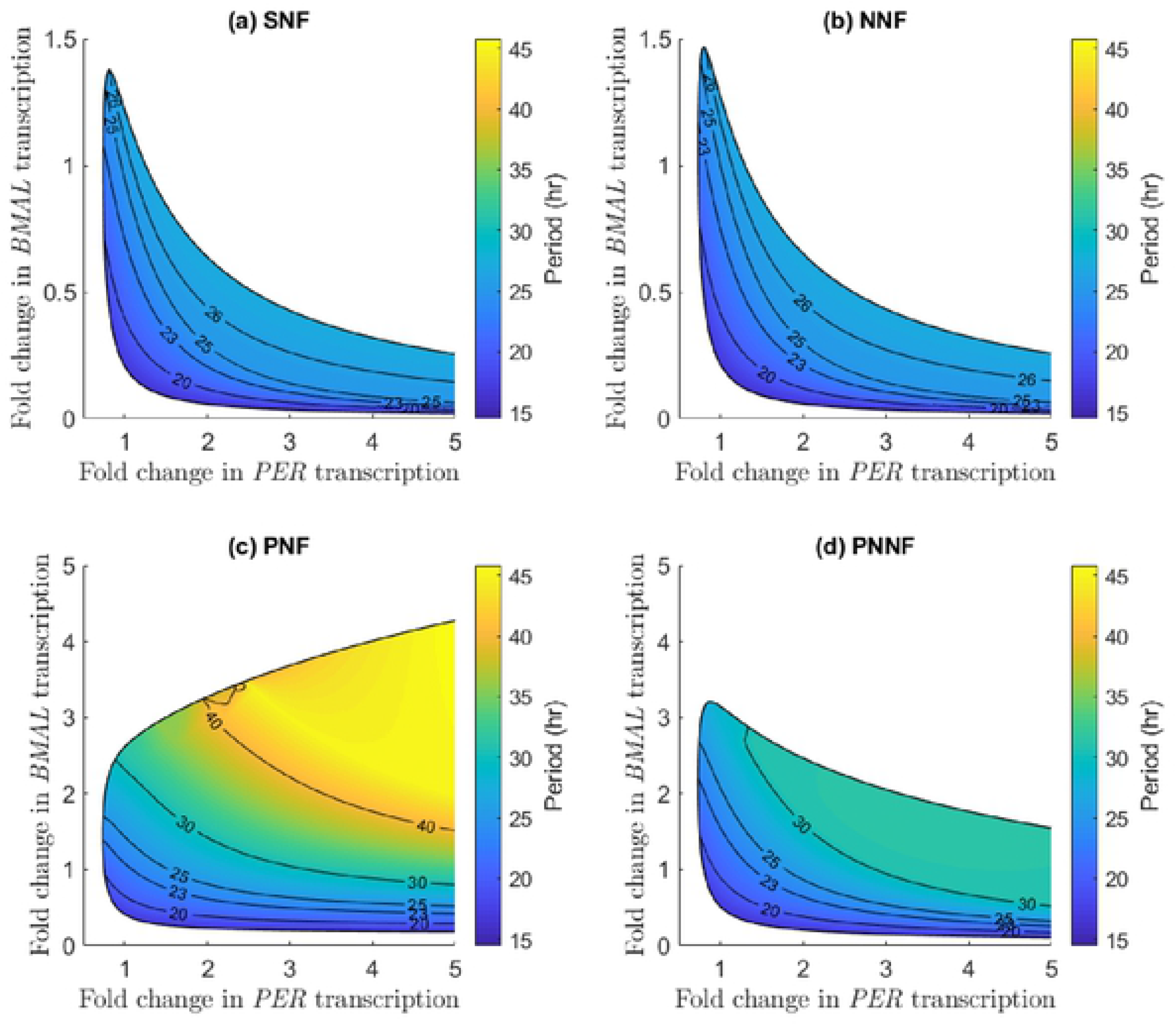
Two-parameter bifurcation curves with respect to fold changes in *BMAL* and *PER* transcription rates for **(a)** SNF(2M8), **(b)** NNF(2M8), **(c)** PNF(2M8), and **(d)** PNNF(2M8). Within the oscillatory regions, period is color-coded. The *BMAL* transcription rate is varied with parameters *A*_T_ (SNF) or *A*_MAX_ (NNF, PNF, PNNF). The *PER* transcription rate is varied with *φ* for all models. All parameter values for these calculations can be found in Table S4.

After our modifications, the models show quite different behavior (Figure 4b). (In comparing Figures 4a and b, keep in mind that *K*_d_ = 10^−5^ in panel a (the KF value) but *K*_d_ = 10^−1^ (more realistic value) in panel b.) In panel b, it is evident that the stoichiometric balance between PER and BMAL required for oscillations in the KF models is not preserved in our models, which is due to the introduction of the Michaelis-Menten rate law for degradation of nuclear PER, as suggested by Figure S2. Furthermore, in our modified models, the NNF and SNF regions of oscillation are approximately equal, implying that the additional negative feedback loop is not having much of an effect on the system’s behavior. Kim and Forger concluded that the additional negative feedback through REV-ERB must be slow compared to the core negative feedback loop in order to provide a robustness benefit to the SNF loop (27). However, in our version of the models, it appears that neither increasing the length of the REV-ERB feedback loop (Figure S3a) nor decreasing the time scaling factor *δ* (Figure S3b) changes the NNF model’s robustness. Thus, in the modified models, the role of the auxiliary negative feedback loop is unclear. Lastly, in our implementation, the PNF model has by far the largest region of oscillations, and the PNNF model shares characteristics of the NNF and PNF models.

### All Four Models (SNF, NNF, PNF and PNNF) Have Comparable Robustness with Respect to ~24 Hour Oscillations

In the previous subsection, we were concerned only with the existence of oscillations and not with their periods. Figure 5 redraws the bifurcation plots in Figure 4b with colors to indicate the oscillatory periods. (To convert from dimensionless period *τ* to period in hours, we assume that *β*_1_ = 1 h^−1^, which makes the period of SNF(2M8) model with its default parameter values (Table S4) around 24 h (Figure S5).) The oscillatory period for each model varies over characteristic ranges. The modified SNF and NNF models have nearly identical period distributions over a range of 16 – 26 h, supporting our conclusion that the additional negative feedback loop has little effect on the system’s periodicity. The PNF and PNNF models have broader period distributions: 15 – 45 h and 16 – 39 h, respectively, confirming the common understanding that a combination of positive and negative feedback loops generates robust oscillations over a broad range of periods (42, 44). Nonetheless, Figure 5 also indicates that all four modified models exhibit similar bands of circadian periodicity (23 – 25 h). So, while the PNF model has a much larger oscillatory domain than the other models, much of this area corresponds to non-circadian periods.

Finally, we note that the PNF(2M8) model exhibits subcritical Hopf bifurcations, which expand the region of long-period (> 40 h) oscillations, as shown in Figure S4.

## CONCLUSION

The Kim-Forger (KF) models of mammalian circadian rhythms (called SNF, NNF and PNF) are appealing in many respects; but they rely on an unrealistic requirement for oscillations, namely that the equilibrium dissociation constant of the PER:CRY::BMAL:CLOCK complex must be *K*_d_ < 10^−3^ nM, which is orders of magnitude smaller than a reasonable value for protein-protein interactions (*K*_d_ ≈ 1—100 nM). This difficulty can be ameliorated (1) by lengthening the core negative feedback loop between *PER* mRNA transcription and PER:CRY inactivation of BMAL:CLOCK (the transcription factor driving *PER* expression), and/or (2) by implementing a Michaelis-Menten rate law for the degradation of nuclear PER. The KF models were further modified by introducing alternative rate laws for BMAL:CLOCK-mediated transcription of clock genes (*PER*, *REV-ERB* and *ROR*) to correct a problem at low expression of the *BMAL* gene, and to provide more accurate rate laws for the effects of REV-ERB and ROR on *BMAL* expression.

With these modifications, we find (Figure 3b) that the SNF model can exhibit oscillations for *K*_d_ as large as 25 nM for 0.05 nM < *A*_T_ < 50 nM. These values are consistent with experimental measurements of *in vitro* binding between PER:CRY and BMAL (41) and of the total number of BMAL and CLOCK molecules in mouse cells (45).

The simple negative feedback loop (SNF), whereby PER inhibits its own synthesis, can be supplemented with an auxiliary positive feedback from ROR (PNF) or a negative feedback from REV-ERB (NNF) on the synthesis of BMAL. A fourth model (PNNF) considers all three feedback loops together. The behaviors of our modified models differ significantly from Kim and Forger’s original conclusions. Kim and Forger’s observation of a stoichiometric balance between PER and BMAL was not observed in the modified models. The modified models also do not share the same trends in robustness (in terms of the size of the oscillatory domain in parameter space). In the original KF models, NNF was the most robust model and PNF the least, but in our modified models, PNF was considerably more robust than the other three models. Nonetheless, the domains of circadian oscillations (23-25 h) are quite comparable for all four of our models. Since the auxiliary positive and negative feedback loops do not seem to provide any additional circadian robustness to the simple negative feedback loop, the functions of ROR and REV-ERB in generating circadian rhythmicity are still unclear in our versions of the KF models.

We propose that these models can be extended in future publications to explore other known behaviors of the mammalian circadian clock. For instance, our models do not address the circadian clock’s temperature-compensation or phase-shifting properties. Adding these key features may answer some remaining questions about the behaviors of these models. For example, the additional negative feedback loop between REV-ERB and BMAL may contribute to these features. Another question that could be addressed with these models is the function of an anti-sense transcript of the PER2 gene (46). Furthermore, these models could be applied in chronotherapy and chrono-pharmacology studies. One such application would be modeling PER2’s interaction with the tumor suppressor protein p53 in stressed (e.g., DNA damage) cells compared to un-stressed cells (47, 48).

## Acknowledgements

This paper is based on a thesis submitted by BH in partial fulfillment of a B.S. degree in Systems Biology from Virginia Tech. JC is supported by NIH (1R35GM138370).

